# Transcriptome Analysis of the Nematode *Caenorhabditis elegans* and *Litoditis marina* in a Chemically Defined Food Environment

**DOI:** 10.1101/2022.01.28.478261

**Authors:** Peiqi Sun, Xuwen Cao, Liusuo Zhang

## Abstract

Diets regulate animal development, reproduction, and lifespan. However, the underlying molecular mechanisms remain elusive. A chemically defined CeMM diet attenuates development and promotes longevity of *C. elegans*, but whether it impact on other nematodes is unknown. Here, we studied the effect of the CeMM diet on the development and longevity of the marine nematode *Litoditis marina*, which belongs to the same family as *C. elegans*. We further investigated genome-wide transcriptional responses to CeMM and OP50 diet for both nematodes, respectively. We observed that the CeMM diet attenuated *L. marina* development but did not extend its lifespan. We found that many of the FOXO DAF-16 target genes, lysosome and xenobiotic metabolism related genes were significantly increased on the CeMM, which might contribute to the lifespan extension of *C. elegans*. Notably, we found that the expression of lysosome and xenobiotic metabolism pathway genes was significantly down regulated in *L. marina* on CeMM, which might explain why the CeMM diet could not promote the lifespan of *L. marina* compared to bacterial feeding. Additionally, down-regulation of several RNA transcription and protein generation and related processes genes might not only be involved in extending longevity but also contribute to attenuating development of *C. elegans* on CeMM, while down-regulation of unsaturated fatty acids synthesis genes might contribute to slow down the growth of *L. marina* on CeMM. Further genetic analysis of candidate gene(s) of longevity and development in *C. elegans* and *L. marina* will provide the molecular mechanisms underlying how diets regulate animal physiology and health in the context of global climate change with variable nutritional environments.

## INTRODUCTION

Diets regulate animal development, reproduction, and lifespan (MacNeil et al., 2013). Different dietary habits can affect the structure and metabolome of gut microbiota and may contribute to health or the pathogenesis of disorders such as atherosclerosis, coronary vascular disease, and inflammatory bowel disease both in humans and mice (Albenberg and Wu, 2014; Lindskog Jonsson et al., 2018). Diets modulate lifespan, consumption, and fat deposition in flies (Skorupa et al., 2008). Dietary restriction, which influences organelle function, gene expression, neural signaling, can extend the lifespan of every species that has been tested, including invertebrates and mammals, and also reduce the incidence of and/or slows the progression of many age-related pathologies (Bishop and Guarente, 2007).

*Caenorhabditis elegans*, as a model organism for biomedical research, has been widely used in nutritional regulation of development and longevity (Szewczyk et al., 2006; Muschiol et al., 2009; Zhang et al., 2010; Pang and Curran, 2014; Green et al., 2022). It was known that *C. elegans* developed from L1 larva to adult in about 3 days by feeding on bacteria *E. coli* OP50, the standard laboratory food, at 20 °C (Brenner, 1974). In *C. elegans*, distinctive food can affect its development, reproduction, and lifespan (Shtonda and Avery, 2006; Coolon et al., 2009; Soukas et al., 2009). For instance, feeding on the bacteria, *Comamonas sp*. led *C. elegans* to lay fewer eggs and exhibit a shorter lifespan compared to the *E. coli* OP50 diet (MacNeil et al., 2013). *C. elegans* maintenance medium (CeMM) was developed as an axenic chemically defined food source, a suboptimal food source for wild-type *C. elegans* (Lu and Goetsch, 1993; Szewczyk et al., 2003). Our previous study showed that only about 10% of the N2 larvae developed into adults until 8 days post-hatching when grown on CeMM; the development is significantly attenuated compared to bacterial feeding (Zhang et al., 2015). In particular, under CeMM condition, this report also showed that the mean life span of *C. elegans* is about 58 days, and the maximum could reach 90 days. In contrast, under OP50, the mean life span of *C. elegans* is 18-20 days (Zhang et al., 2020). Due to exclude variables associated with bacterial metabolism (Dougherty and Calhoun, 1948; Szewczyk et al., 2003), the CeMM diet is a powerful platform to study gene by environment interactions, pharmaceutical drug screening, and space biology in combined with *C. elegans* (Szewczyk et al., 2006; Zhang et al., 2015; Cao et al., 2021).

After five years of intensive efforts, our research group has developed *L. marina* as a promising marine nematode model, which could be used as a marine satellite animal model to the terrestrial biomedical model animal *C. elegans* (Cong et al., 2020; Xie et al., 2020; Xie et al., 2021; Zhao et al., 2021). Given that *L. marina* and *C. elegans* belong to the same family of *Rhabditidae* in the *phylum* Nematoda (Xie et al., 2020), it allows us to compare the physiological changes of marine and terrestrial relatives in response to changing nutritional environments. Our previous reports showed that newly hatched L1 larvae reach sexual maturity within 4-5 days when feeding bacteria *E. coli* OP50 (Xie et al., 2020), and the life span of *L. marina* is comparable to *C. elegans* in the bacterial diet (Xie and Zhang, 2021).

In this report, we found that CeMM diets attenuated the development of *L. marina*, compared to feeding bacteria, which is in line with *C. elegans*. However, we observed that the lifespan of *L. marina* on the CeMM diet was comparable to that of bacterial feeding, in contrast to the previous report that the CeMM diet promotes the longevity of *C. elegans*. We further investigated their transcriptomic signatures via RNA sequencing (RNA-seq) in different food environments. We found that many of the *daf-16* target genes and genes involved in the lysosome and xenobiotic metabolism were significantly increased on the CeMM diet, which might contribute to the lifespan extension of *C. elegans*. In addition, we found that the expression of protein synthesis-related genes was significantly down-regulated on CeMM, which might also promote the longevity of *C. elegans* on the CeMM diet. Notably, we found that the expression of lysosome and xenobiotic metabolism pathway genes was significantly down-regulated in *L. marina* on CeMM, which might explain why the CeMM diet could not promote the lifespan of *L. marina* compared to bacterial feeding. We found that the expression of genes in several pathways, such as tryptophan metabolism and fatty acid degradation, was significantly increased in both *C. elegans* and *L. marina* on CeMM compared to bacterial feeding, which might contribute to the developmental attenuation for both nematodes on CeMM. In addition, down-regulation of several RNA transcription and protein generation and related processes genes might attenuate the development of *C. elegans*, while down-regulation of unsaturated fatty acids synthesis genes in *L. marina* may contribute to its slow development on CeMM. Further genetic analysis of the key candidate gene(s) will provide the molecular mechanisms underlying how diets regulate animal physiology and health in the context of global climate change with variable nutritional environments.

## MATERIALS AND METHODS

### Cultivation of strains

The *C. elegans* wild-type Bristol N2 strain was obtained from the Caenorhabditis Genetics Center (CGC). At 20 *°C*, OP50 bacteria were used to culture the *C. elegans* strain on NGM plates using standard methods (Brenner, 1974). For the CeMM part, the 1.7% agarose CeMM plate was used directly at 20 °C when conducting worm growth observation (Szewczyk et al., 2006). *L. marina* was grown at 20 °C on seawater NGM plates inoculated with *E. coli* OP50. 15‰ sea salt CeMM-agarose plates were prepared for *L. marina* experiments.

### Worm Synchronization

To obtain synchronized nematodes, the hermaphrodites of *C. elegans* with full eggs were bleached with basic hypochlorite solution at room temperature until each adult worm was digested (da Silva, 2005). Synchronized L1s were obtained, followed by eggs collection and hatching overnight on unseeded nematode growth medium NGM plates at 20 °C for about 12 hs. Then, *L. marina* were cultivated on a medium of 9 cm. When there were many eggs on the plate, the eggs were collected into a 15 mL centrifuge tube at 1300 g for 1 min. After being washed twice with sterilized water, the eggs were treated with alkaline hypochlorite solution at room temperature, 60-90 s, 1300 g for 1 min. The egg liquid was incubated in sterilized seawater for 21 h. After incubation, the unhatched eggs and remnants were removed by 450 mesh filtrations with sterilized seawater after extraction. Then 500 mesh filter was for a second filtration. After centrifugation for 1min, the synchronized L1s were obtained.

### CeMM preparation

2×CeMM stock media was prepared with determinate chemicals, including vitamins and growth factor solutions, salt solutions, amino acids, nucleic acid substituents (Cao et al., 2021). The above chemical solutions were mixed, and the mixed media was filter sterilized using a Corning 0.2 mm filter system, and the final stock media were stored at 4 °C. When using, 100 mL 2×CeMM final stock media were mixed with 100 mL 3.4% agarose, diluting the final CeMM content 1× and agarose content 1.7%. The nystatin (200 mg nystatin/14 mL 95% ethanol) was subsequently added to the mixed solution. For 15‰ sea salt CEMM-agarose, 3 g sea salt were added when prepared 3.4% agarose. All operations are performed in a sterile environment. Divide the mixed media into Petri dishes of 3.5 cm diameter, which contained 3 ml of CeMM-agarose. The CeMM-agarose plate was kept at 4 degrees away from light directly.

### CeMM growth and lifespan Assay

Seventy synchronized newly hatched L1s of *L. marina* were transferred and cultured in 3-cm-diameter CeMM plates. The growth and development of worms were observed, and the adults were counted and picked out every day. Three replicates were performed for each strain. For lifespan assay, 40-60 L4 females were transferred to 15‰ sea salt CEMM-agarose plates. The number of live and dead worms was determined using a dissecting microscope every 48 h. Live worms were transferred to fresh 15‰ sea salt CEMM-agarose plates. Worms were scored dead if no response was detected after prodding with a platinum wire. Dead worms on the wall of the plate were not counted. Three replicates were performed for each strain.

### RNA-seq library preparation and RNA-seq analysis

To acquire synchronized *C. elegans* newly hatched L1 larvae, the adults with full eggs were bleached using standard hypochlorite to obtain embryos. Larvae were hatched and underwent growth arrest on unseeded plates (da Silva, 2005). Synchronized L1 larvae of N2 were transferred to 90 mm NGM with full of *E. Coil* and CeMM plates with treating 2.5 hours, respectively. Similarly, synchronized L1s of *L. marina* were transferred to 90 mm Sea-Water (SW) plates full of *E. Coil* and 15‰ sea salt CEMM-agarose plates for feeding 2.5 hours, respectively. Three replicates were performed both for N2 and *L. marina*, respectively. The worms were washed with M9 three times to remove remnants of the medium. The samples were then transferred to 1.5 ml tubes, and the excess supernatants were removed via centrifugation (1,300 g, 1 min). Remove the liquid samples as soon as possible and then freeze them immediately in liquid nitrogen for 5 minutes and store them at −80 °C. RNA was extracted using the Trizol method after samples that were swiftly ground with liquid nitrogen. A total amount of 1 μg RNA per sample was used as input material for the RNA sample preparations. After reverse transcription, cDNA fragments of preferentially 370~420 bp in length were selected and purified to prepare for the library. After the library was constructed, Qubit3.0 Fluorometer was used for initial quantification. Nanodrop 2000 and Agilent 2100 BioAnalyzer were then used to detect the insert size of the library, and qRT-PCR accurately quantified effective library concentration to ensure library quality. Followed by a qualified library inspection, Illumina sequencing was performed, and 150 bp paired-end reads were generated. Reference genome and gene model annotation files were downloaded from Ensembl (*Caenorhabditis elegans* (WBcel235)). The reference genome index was built by Hisat2 (Kim et al., 2015), and paired-end clean reads were aligned and counted to the reference genome using StringTie (Pertea et al., 2015). And then, FPKM was calculated based on the length of the gene and the reads count mapped for each gene (Florea et al., 2013). Differential expression analysis of two groups (two biological replicates per condition) was performed using the DESeq2 R package (1.20.0) (Love et al., 2014). Use Benjamini and Hochberg’s approach to control the false discovery rate for the resulting P-values. Correcting P< 0.01 and fold change> 2 were used as the threshold for significantly differential expression. GO enrichment analysis and KEGG pathway enrichment analysis of differentially expressed genes were realized by clusterProfiler R package (Yu et al., 2012).

### *daf-16* target genes identification

According to the references (Li and Zhang, 2016; Sun et al., 2017), a total of 118 *daf-16* direct targets genes were included in our analysis, among them 109 were from Li’s report and 22 genes were from in Sun’s paper, and 13 genes are overlapping in the two reference. These DAF-16 target genes were compared with our RNA-seq data by ‘WormBase_ID’ using Micro Office Excel. As a result, 41 DAF-16 target genes were identified in *C. elegans* on CeMM, 11 of which were enriched in our KEEG analysis.

### Real-time quantitative PCR (qPCR)

Some of the critical genes of our interest were selected for qPCR validation; glutathione S-transferase gene *gst-4*, acyl-coenzyme A oxidase gene *acox-3*, superoxide dismutase gene *sod-3*, eukaryotic initiation factor gene *eif-6*, heat shock protein gene *hsp-1 of C. elegans*, glutathione S-transferase gene EVM0002486/*gst-8*, dehydrogenases short-chain gene EVM0002754/*dhs-2*, UDP-glucuronosyl transferase EVM0005671/*ugt-19*, fatty acid elongation EVM0013887/*elo-3 of L. marina*.

Synchronized L1 worms of *C. elegans* and *L. marina* were separately grown on OP50 with NGM (or SW-NGM) and CeMM (or 15‰ sea salt CEMM) at 20 °C for 2.5 h. Quantitative PCR total RNA was isolated using Trizol (Invitrogen), and standard cDNA synthesis was performed using a cDNA synthesis kit (Toyobo, FSQ-301). We performed reverse transcription with 500 ng RNA per sample. PCR amplification, using SYBR Green (Toyobo, QPK-201), via ABI QuantStudio 6 Flex system, was performed in 96-well microtiter plates. The internal reference gene reflected the gene expression. Gene names and Primers used for real-time PCR were shown in Supplementary File 1. Quantitative PCRs (qPCRs) were performed in triplicate.

## RESULTS

### CeMM Diet Attenuates the Development of *L. marina* While Does not Extend Its Longevity

To ask whether the CeMM diet affects the development of *L. marina*, we transferred newly hatched L1 larvae into 15‰ sea salt CEMM and found that only about 15% of them developed into adulthood after 8 days post-hatching when grown on CeMM (Fig 1A). To ask whether the CeMM diet promotes the lifespan of *L. marina* compared to bacterial feeding, we transferred L4 female larvae to 15‰ SW-CeMM plates. We observed that *L. marina* could live as long as 28 days with an average lifespan of about 16 days (Figure 1B), which is similar to that of OP50 feeding (Xie and Zhang, 2021). The data suggested that the CeMM diet attenuated the development of *L. marina* but did not promote its longevity compared to bacterial feeding.

**FIGURE 1.**
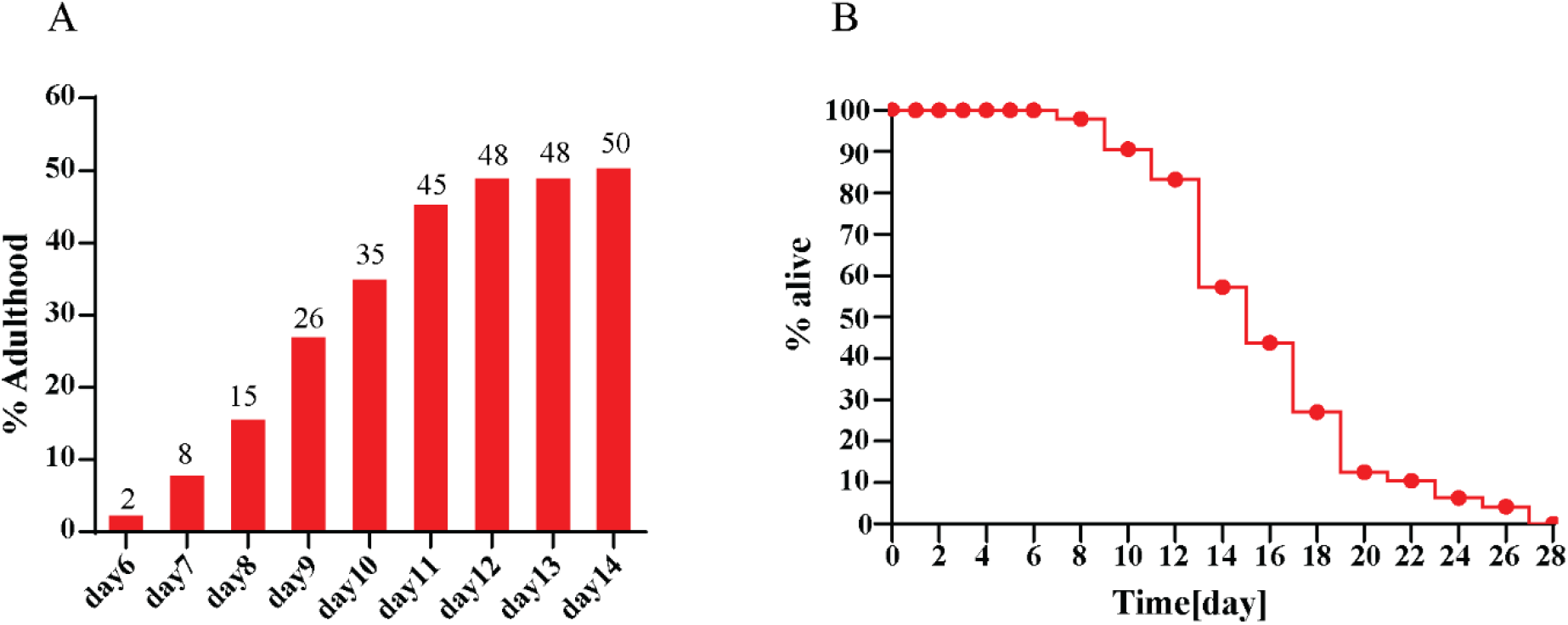
CeMM diet attenuates the development of *L. marina* while does not promote its longevity. (A) For developmental analysis, 70 newly hatched L1s were transferred onto each indicated 3 cm-dimeter agar plates15‰ CeMM. Three replicates were performed. (B) Lifespan assay, 55 L4 females were transferred to each assay plate, incubated at 20°C. The number of live and dead worms was counted every day. Live worms were assigned to 3 cm-dimeter agar plates15‰ sea salt CeMM plates every two days. Three replicates were conducted.

### RNA-Seq Analysis in *C. elegans* and *L. marina* under CeMM Food Environment

To investigate the genome-wide transcriptional responses of *C. elegans* and *L. marina* in CeMM and OP50 dietary conditions, we used RNA-seq analysis to describe their transcriptomic signature. Both newly hatched L1s of *C. elegans* and *L. marina* were treated 2.5 h on CeMM and OP50 plates, respectively. We used OP50 as the control group, CeMM as the treatment group, and FDR < 1% and fold change > 2 were applied as the differential gene screening threshold. A total of 2,571 DEGs for *C. elegans* and 678 DEGs for *L. marina* were identified in our analysis (Fig 2). Details of significantly up and down-regulated DEGs were listed in Supplementary File 2.

**FIGURE 2.**
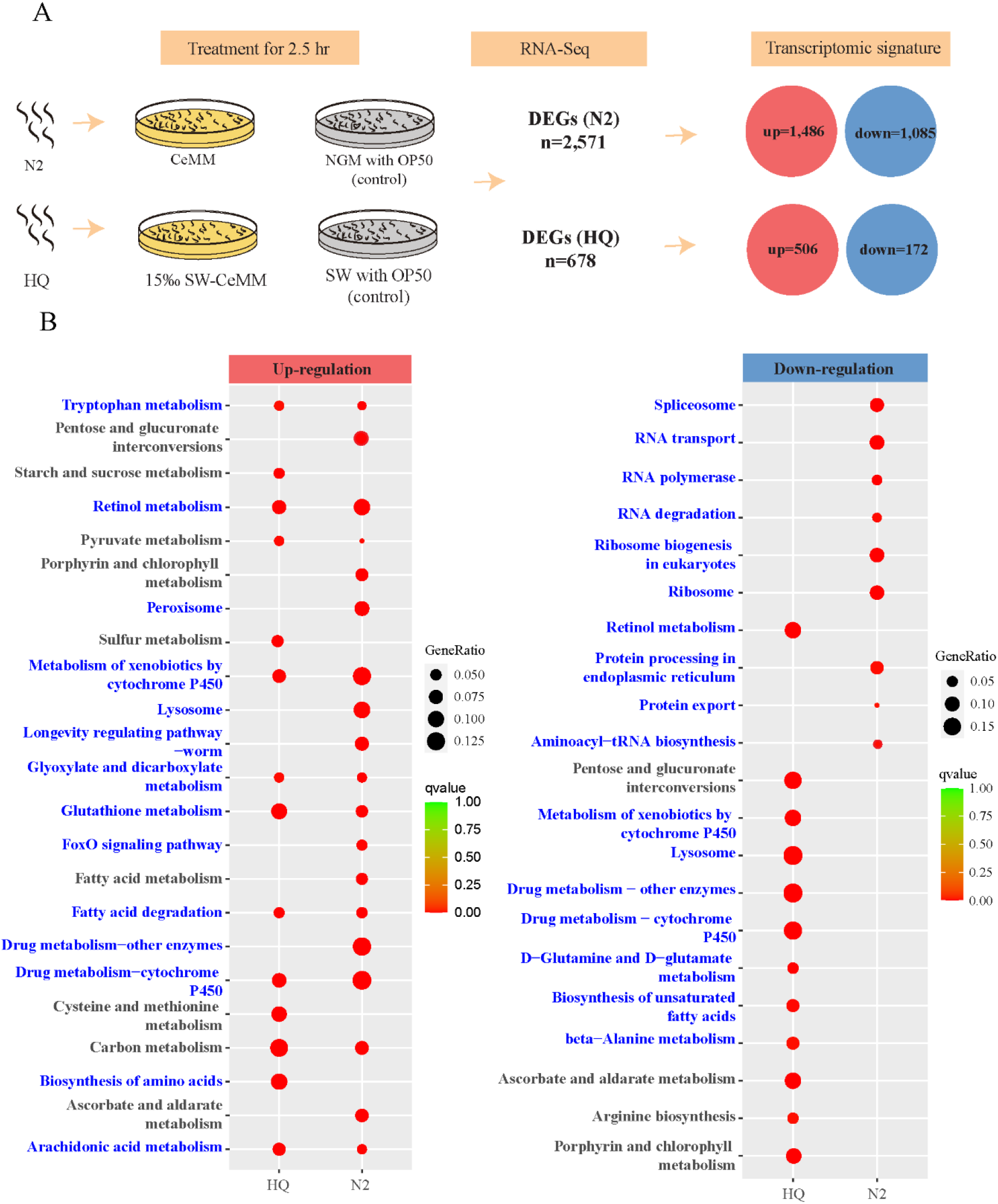
The transcriptomic signature of *C. elegans* and *L. marina*. (A) Experimental design of this study and the resulting transcriptomic signature of worms. Differentially expressed genes (DEGs, foldchange >2; DESeq2 padj < 0.01) were determined for each condition. (B) KEGG enrichment analysis for DEGs. ‘HQ’ and ‘N2’ represent *L. marina* and *C. elegans*, respectively. ‘Up-regulation’ indicates increased gene expression in CeMM diet compared to OP50 diet. ‘Down-regulation’ indicates decreased gene expression in CeMM diet compared to OP50 diet. foldchange >2; DESeq2 padj < 0.01set as the differential gene screening threshold. The color from red to green represents the significance of the enrichment. We focus on the pathway of blue font enrichment. GeneRatio was defined as the ratio of the number of differential genes annotated to the GO term to the total number of differential genes.

### Changed Transcriptomic Signature of *C. elegans* under CeMM Conditions

Based on KEGG analysis, we found that the longevity regulating pathway genes were significantly up-regulated on CeMM, including seven *daf-16* target genes (*mtl-1*, *mlt-2*, *ctl-1*, *ctl-2*, *ctl-3*, *lgg-1*, and *sod-3*) (Fig 3A). In addition, we found that the forkhead box (FOXO) signaling pathway genes were also significantly up-regulated on CeMM (Figure 3B), which include nine potential *daf-16* target genes. In addition to the five *daf-16* target genes shared in the longevity regulation pathway, four additional genes are *mfb-1*, *dct-1*, *aakg-4* and *pck-1*. The worm FOXO orthologue, DAF-16, activates a program of genes to extend longevity, promote resistance to the environment and regulate growth (Arden, 2008). Among these enriched target genes, *lgg-1* has been identified as the *daf-16* direct target involved in longevity (Sun et al., 2017). Furthermore, we found that Lysosome (Figure 3C) and Peroxisome (Figure 3D) related genes were significantly up-regulated under CeMM food conditions, compared to bacterial feeding. The lysosome is a membrane-bound cell organelle containing digestive enzymes involved with various cell processes like programmed cell death or apoptosis, and functions as a ‘signaling hub’ to integrate metabolic inputs, organelle interactions, and the control of longevity (Saftig and Klumperman, 2009; Savini et al., 2019). For example, in *C. elegans*, the lysosomal acid lipase LIPL-4 is highly induced under conditions connected to extended lifespan (Lapierre et al., 2011). Peroxisomes support key metabolic platforms for the oxidation of various fatty acids, regulate redox conditions, and play crucial roles in the biosynthesis of essential lipid molecules and longevity (Weir et al., 2017; Jo et al., 2020). For instance, dietary restriction and AMPK increased lifespan require coordination between mitochondria and peroxisomes (Weir et al., 2017). Peroxisomal catalase gene *ctl-2*, which function as an antioxidant enzyme that protects *C. elegans* cells from reactive oxygen species, contribute to the extended lifespan of *C. elegans* (Petriv and Rachubinski, 2004; Chávez et al., 2007; Wu et al., 2012). Of note, *ctl-2* is a DAF-16 target gene. Our data suggested that *daf-16* target genes, lysosome and peroxisome related genes might play essential roles in promoting the longevity of *C. elegans* when feeding the CeMM diet.

**FIGURE 3.**
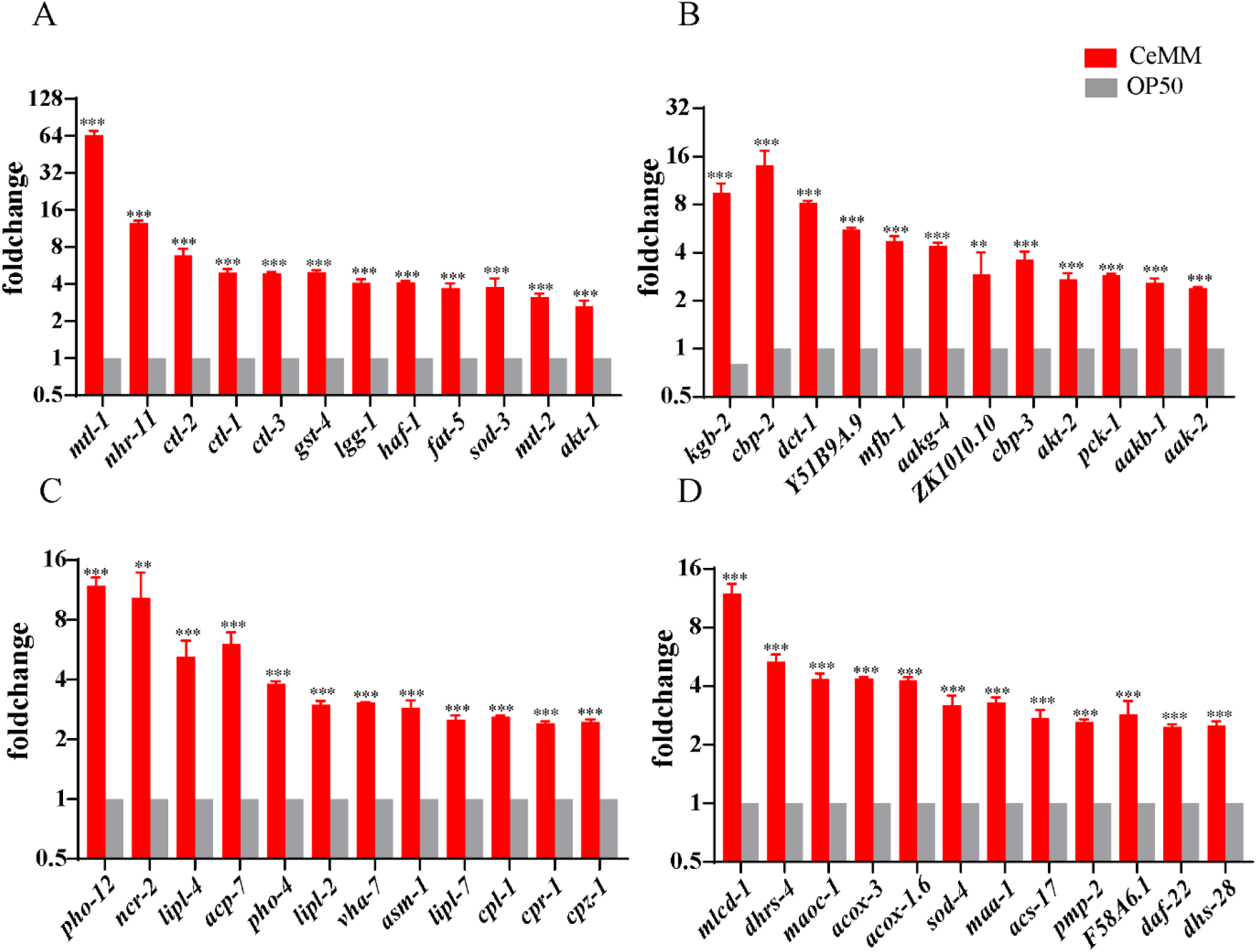
The expression of up-regulated pathway related genes of *C. elegans* on CeMM. (A) The transcript level of longevity regulating pathway related genes. (B) The transcript level of FOXO signaling pathway-related genes. (C) The transcript level of lysosome related genes. (D) The transcript level of peroxisome related genes. Fold changes indicate the ratio of the treatment group (CeMM) to the control group (OP50). The error bars represent the standard error of the mean of three biological replicates per condition. *P < 0.05, **P < 0.01, ***P < 0.001.

We found that ribosome biogenesis, ribosome, RNA transcription and protein generation, processing and transport related genes were significantly decreased in *C. elegans* on CeMM (Fig 4). The expression level of ribosome biogenesis genes such as *eif-6*, *fib-1*, *mac-1*, *nxt-1*, *ran-1* (Fig 4A), ribosome machinery associated genes such as *rpl*-, *rps*-, and *mrpl*-, *mrps*-genes (Fig 4B), and RNA polymerase genes such as *rpc-1*, *rpb-6*, *rpoa-12*, *rpoa-2*, *rpb-12* (Fig 4C), were significantly down-regulated. Ribosomes are macromolecular machines to support biological protein synthesis (Shirokikh, 2021), and RNA polymerase transcripts DNA into RNA such as mRNA, tRNA, rRNA among others (Sainsbury et al., 2015). The down-regulation of these transcription and translation related genes would negatively affect RNA and protein generation and functions. Similarly, spliceosome related genes (*eftu-2*, *emb-4*, *isy-1*, *mog-2*, *snr-1*, Fig 4D), RNA transport-related genes (such as *eif-3. B~eif-3. I*, *hoe-1*, *ife-2*, Figure 4E), Aminoacyl-tRNA biosynthesis related genes (such as *ears-1*, *fars-1*, *fars-3*, *nars-1*, *pars-1*, *tars-1*, *wars-1*, Figure 4F), protein processing in endoplasm genes (*dad-1*, *hsp-90*, *dnj-12*, *hsp-1*, *hsp-17*, Figure 4G), as well as protein export associated genes (such as *emo-1*, *hsp-3*, *oxa-1*, *spcs-2*, *sec-61*, Figure 4H) were significantly down-regulated in response to CeMM diet. Our data suggest that CeMM diet might attenuate development and promote lifespan extension of *C. elegans* through down-regulation of several RNA transcription and protein generation and related processes.

**FIGURE 4.**
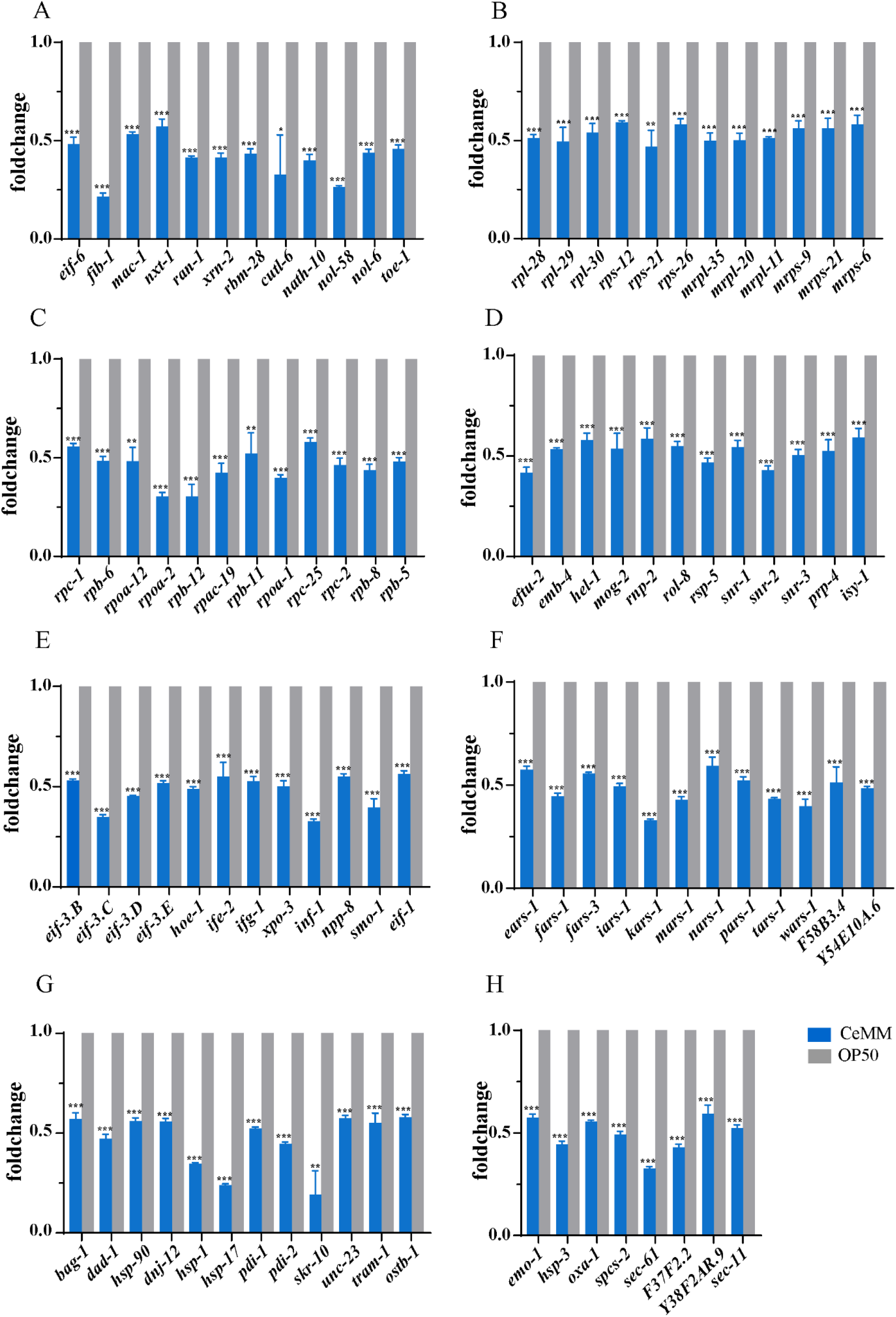
The expression of down-regulated pathway related genes of *C. elegans* on CeMM. (A) The transcript level of ribosome biogenesis in eukaryotes related genes. (B) The transcript level of ribosome genes. (C) The transcript level of RNA polymerase genes. (D) The transcript level of spliceosome related gene. (E) The transcript level of RNA transport related gene. (F) The transcript level of aminoacyl-tRNA biosynthesis related genes. (G) The transcript level of protein processing in endoplasmic related genes. (H) The transcript level of protein export related genes. Fold changes indicate the ratio of the treatment group (CeMM) to the control group (OP50). The error bars represent the standard error of the mean of three biological replicates per condition. *P < 0.05, **P < 0.01, ***P < 0.001.

### Changed Transcriptomic Signature of *L. marina* under CeMM Conditions

In *L. marina*, 678 DEGs were observed under CeMM food conditions compared with bacterial OP50 feeding (Figure 1). We found that three drug and cytochrome P450 pathway genes such as *ugt-19*, *ugt-51*, *ugt-22*, *ugt-23* were significantly down-regulated in *L. marina* on CeMM (Figure 5A). In contrast to *C. elegans*, we found that ‘lysosome’ related genes such as *slc-17.9*, *ptr-22*, *lcn-3.3*, *crn-6* were significantly down-regulated on CeMM (Figure 5B).

**FIGURE 5.**
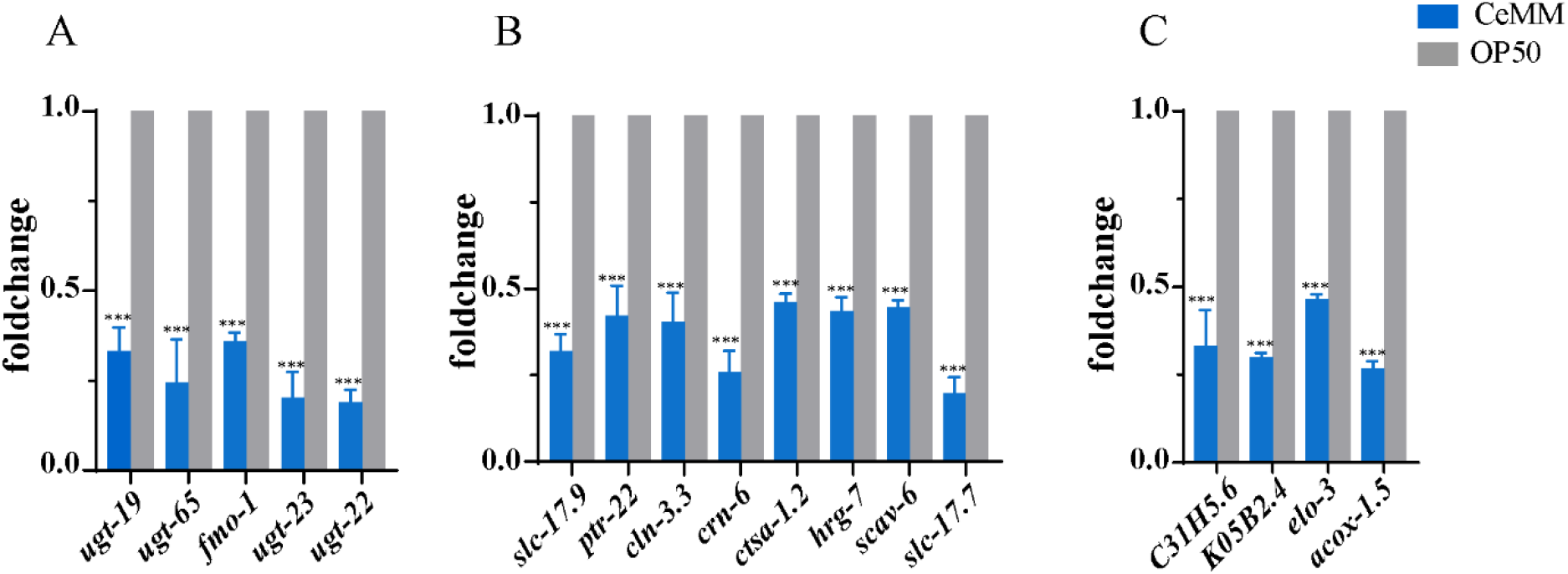
Expression of down-regulated genes of *L. marina* on CeMM. (A) The transcript level of cytochrome of cytochrome P450 pathway related genes. (B) The transcript level of lysosome related genes. (C) The transcript level of Biosynthesis of unsaturated fatty acid related genes. Fold changes indicate the ratio of the treatment group (CeMM) to the control group (OP50). The error bars represent the standard error of the mean of three biological replicates per condition. *P < 0.05, **P < 0.01, ***P < 0.001.

In addition, we found that biosynthesis of unsaturated fatty acid genes such as *elo-3*, *acox-1.5* were also be enriched in *L. marina* down-regulation set (Figure 5C). A previous study showed that mutants with severe unsaturated fatty acid deficiencies display growth and neurological defects in *C. elegans* (Watts, 2002). Our data suggest that CeMM diet might attenuate development of *L. marina* through down-regulation of unsaturated fatty acid synthesis genes.

### Shared Transcriptomic Signature of *L. marina* and *C. elegans* under CeMM Food Condition

We found that the expression of several pathways such as cytochrome P450 (Figure 6A, Supplementary Figure 2A), glutathione metabolism (Figure 2B, Supplementary Figure 2B), retinol metabolism (Figure 6C, Supplementary Figure 2C), carbon metabolism (Figure 6D, Supplementary Figure 2D), arachidonic acid metabolism (Figure 6E, Supplementary Figure 2E), glyoxylate and dicarboxylate metabolism (Figure 6F, Supplementary Figure 2F), tryptophan metabolism (Figure 6G, Supplementary Figure 2G) and fatty acid degradation (Figure 6H, Supplementary Figure 2H) pathway related genes, were significantly increased in both *L. marina* and *C. elegans* on CeMM medium (Figure 1). These shared features indicate that these genes may play essential roles in attenuating development of both *L. marina* and *C. elegans* on CeMM food environments.

**FIGURE 6.**
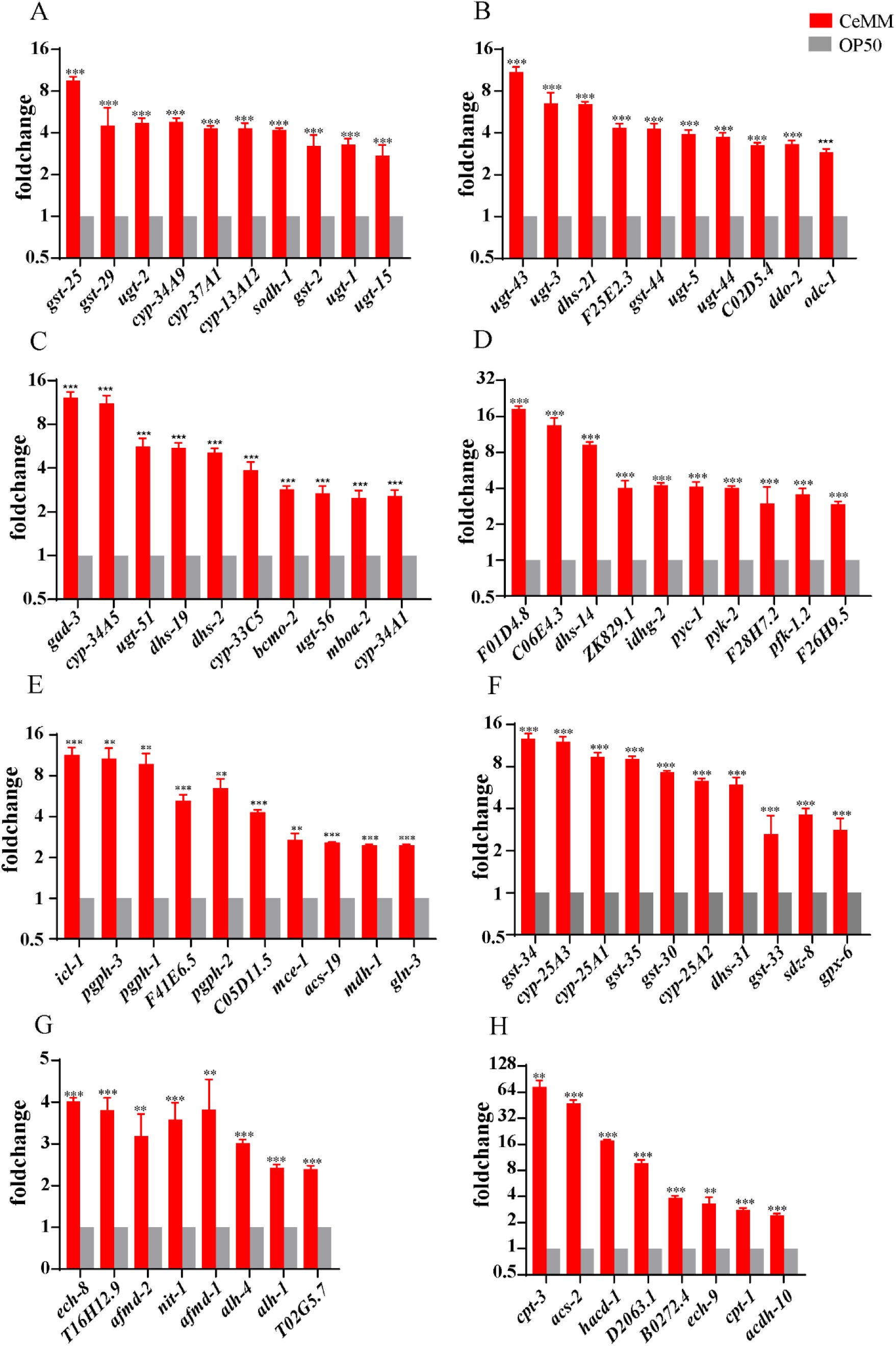
Expression of shared pathway related genes in *C. elegans* on CeMM. (A) The transcript level of cytochrome P450 related genes in *C. elegans*. (B) The transcript level of glutathione metabolism related genes in *C. elegans*. (C) The transcript level of retinol metabolism genes in *C. elegans*. (D) The transcript level of carbon metabolism genes in *C. elegans*. (E) The transcript level of glyoxylate and dicarboxylate metabolism related genes in *C. elegans*. (F) The transcript level of arachidonic acid metabolism related genes in *C. elegans*. (G) The transcript level of Tryptophan metabolism related genes in *C. elegans*. (H) The transcript level of Fatty acid degradation related genes in *C. elegans*. Fold changes indicate the ratio of the treatment group (CeMM) to the control group (OP50). The error bars represent the standard error of the mean of three biological replicates per condition. *P < 0.05, **P < 0.01, ***P < 0.001.

### Quantitative Real-Time PCR Validation

We conducted qPCR to testify the expression of interest genes identified from our RNA-seq results (Figure 7A, Supplementary File 5). Consistent trends were demonstrated in Figure 7B. The expression of glutathione S-Transferase gene *gst-4*, acyl-Coenzyme A oxidase gene *acox-3*, superoxide dismutase gene *sod-3* of *C. elegans*, and glutathione S-Transferase gene EVM0002486/ *gst-8*, dehydrogenases short chain gene EVM0002754/*dhs-2* were significantly up-regulation when growing on CeMM. On the contrary, the expression of eukaryotic initiation factor gene *eif-6*, heat shock protein gene *hsp-1* of *C. elegans*, UDP-Glucuronoxylan transferase *EVM0005671/ugt-19*, fatty acid elongation EVM0013887/*elo-3* were significantly down-regulation under the CeMM diet.

**FIGURE 7.**
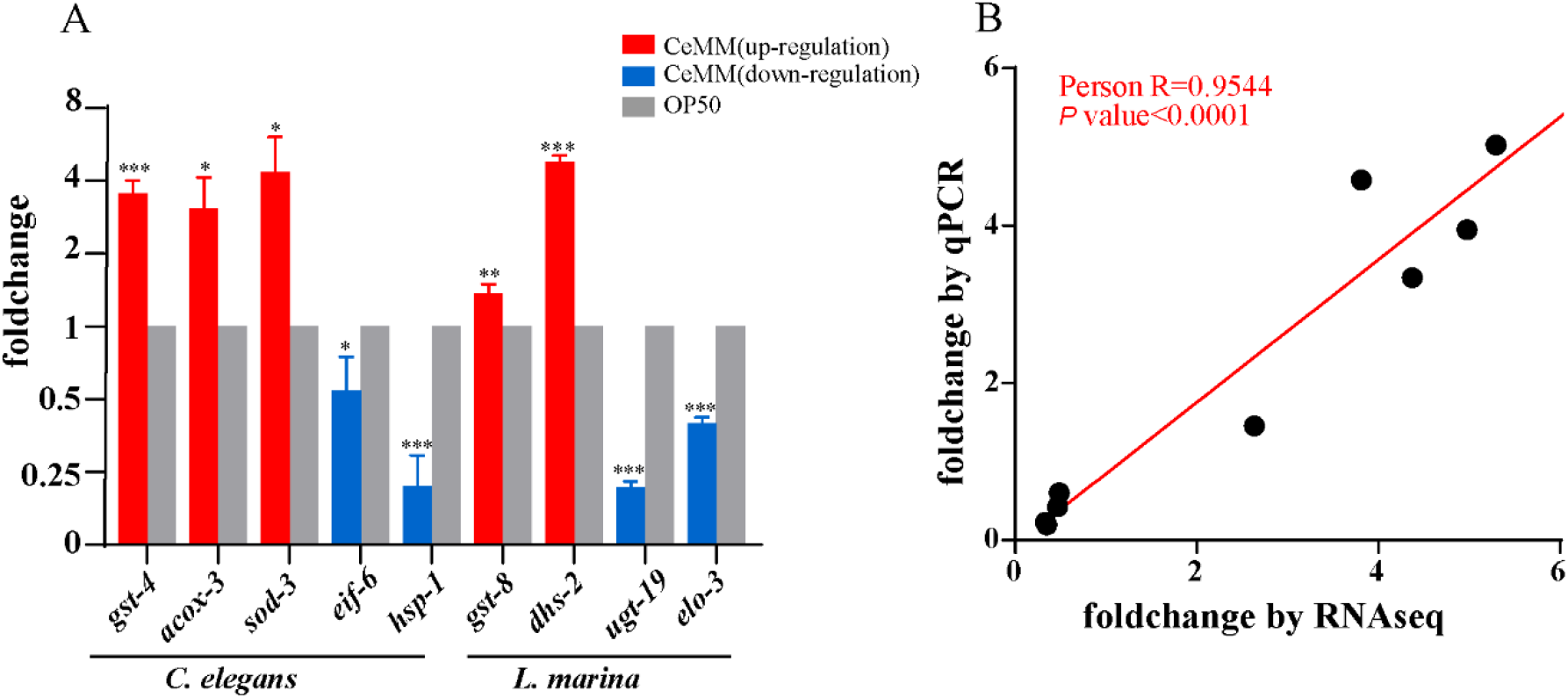
Validation of the RNA-seq results using qPCR. (A) qPCR analysis of interest genes identified from the RNA-seq results. Fold changes indicate the ratio of the treatment group (CeMM) to the control group (OP50). The error bars represent the standard error of the mean of three biological replicates per condition. *P < 0.05, **P < 0.01, ***P < 0.001. (B) Correlation analysis of the results of RNA-seq and qPCR for interest genes. Each dot represents a gene, with detailed information shown in Supplementary File 5. Correlation analysis was performed and graphed using GraphPad Prism 8software, Pearson R = 0.9544, with a P-value < 0.0001.

## DISCUSSION

*C. elegans* develop from L1 larvae to adults in about 3 days when fed OP50 at 20 °C (Brenner, 1974), whereas only about 10% of the larvae developed into adults after 8 days post-hatching when grown on CeMM (Zhang et al., 2015). A previous report showed that the mean life span of *C. elegans* is about 16 days when fed on OP50 in 20°C, and the maximum could reach 31 days (Szewczyk et al., 2006), while under CeMM condition, the mean life span of *C. elegans* is about 30 days, and the maximum could go to unexpected 93 days (Szewczyk et al., 2006). Similarly, we previously reported that the mean life span of *C. elegans* is about 58 days, and the maximum could reach to about 90 days (Zhang et al., 2015). The CeMM diet attenuates *C. elegans* development and extends its lifespan (Szewczyk et al., 2006; Zhang et al., 2015). It was reported that *daf-16* and its potential target genes such as *mtl-1* and *mtl-2* are up-regulated in CeMM food conditions (Szewczyk et al., 2006). However, the precise mechanisms underlying how the CeMM diet attenuates the development and promotes the longevity extension of *C. elegans* are largely unexplored, and whether the CeMM diet regulates the growth and lifespan of other nematodes is unknown. We recently showed that around 91% of newly hatched L1 larvae of *L. marina* developed into adulthood after 5 days at 20°C, and the mean life span of *L. marina* is about 16 days (Xie and Zhang, 2021; Xie et al., 2021). In this report, we observed that the CeMM diet attenuated the development of the marine nematode *L. marina* while not promoting its lifespan. Previous microarray analysis identified 48 significantly altered genes (Szewczyk et al., 2006). We compared these 48 genes to our DEG data and found that 12 of which were significantly enriched in *C. elegans* on CeMM, six of them were expressed in the same trend and the expression of another six genes was in the opposite direction (Supplementary File 4). The six overlapping genes changing in the same direction include *mtl-1*, *mtl-2*, *gale-1*, *zhit-3*, *K05C4.5* and *dnj-10*. Among them, *mtl-1* and *mtl-2* were significantly increased, while *gale-1*, *zhit-3*, *K05C4.5* and *dnj-10* were significantly reduced on the CeMM diet. Notably, *mtl-1* and *mtl-2* were enriched in the “longevity regulating pathway” KEEG terms in *C. elegans* (Figure 3A), and both of them are FOXO DAF-16 target genes. It is deserved to further study the function of these overlapping genes in longevity regulation.

### CeMM Promotes the Longevity of *C. elegans* but not *L. marina*

FOXO transcription factor DAF-16 and its target genes play essential roles in aging (McElwee et al., 2003; Murphy et al., 2003; Dong et al., 2007; Schuster et al., 2010; Li and Zhang, 2016; Sun et al., 2017). We found that 11 potential *daf-16* target genes such as *mtl-1*, *mtl-2*, *ctl-1*, *ctl-2*, *ctl-3*, *lgg-1*, *sod-3*, *mfb-1*, *dct-1*, *aakg-4*, *pck-1* were enriched in *C. elegans* on CeMM (Figure3 A, B). RNAi of the metallothionein *mtl-1, the cytosolic and* peroxisomal catalases *ctl-1* and *ctl-2* and the mitochondrial superoxide dismutase *sod-3*, repaired the lifespan-extending properties of active DAF-16 (Murphy et al., 2003), and RNAi of *lgg-1* suppresses the increased lifespan induced by LIPL-4 overexpression (O’Rourke et al., 2013). Further, in addition to the 11 *daf-16* target genes enriched in our KEEG analysis (Figure 3 A, B), we found another 30 DAF-16 direct targets according to the previous studies (McElwee et al., 2003; Murphy et al., 2003; Dong et al., 2007; Schuster et al., 2010; Li and Zhang, 2016; Sun et al., 2017). Among them, 22 genes were significantly increased, and eight were significantly down-regulated in *C. elegans* on CeMM (Supplementary File 3). Previous studies showed that the inactivation of the L-malate dehydrogenase gene *mdh-1* could significantly reduce the *daf-2* mutant lifespan (Samuelson et al., 2007). Of note, some of the *daf-16* targets such as *maple-12*, *sams-1*, and *ubh-4* were anti-longevity genes, and RNAi knock-down *mrpl-12*, which encodes a mitochondrial ribosomal protein, *sams-1*, which encodes S-adenosyl methionine synthetase, and *ubh-4*, which encodes ubiquitin C-terminal hydrolase, could extend lifespan of *C. elegans*, respectively (Hamilton et al., 2005; Hansen et al., 2005; Kim and Sun, 2007; Matilainen et al., 2013). The expression of *mrpl-12, sams-1*, and *ubh-4* was significantly repressed in *C. elegans* on CeMM. Our data suggested that the CeMM diet might extend the longevity of *C. elegans* on CeMM via FOXO *daf-16* signaling. Notably, the longevity and FOXO pathway related genes were not enriched in *L. marina* on the CeMM diet.

Lysosome has been reported as a ‘signaling hub’ that integrate metabolic inputs, organelle interactions, and the control of lifespan in *C. elegans* (O’Rourke and Ruvkun, 2013; Settembre et al., 2013a; Settembre et al., 2013b; Savini et al., 2019; Sun et al., 2020). Lysosomal acid lipase (*lipl*) has been found to increase autophagy, which extends longevity in *C. elegans* and mouse autophagy (Lapierre et al., 2011; Palmieri et al., 2011), and overexpression of LIPL-4 significantly increase the lifespan of *C. elegans* (Wang et al., 2008; Folick et al., 2015). In this report, three *lipl* genes, *lipl-2*, *lipl-7*, *lipl-4* were significantly up-regulated in *C. elegans* on CeMM (Fig 3C). Our data suggested that the CeMM diet might promote the longevity of *C. elegan* on CeMM via lysosome related functions. Of note, the expression of lysosome related genes was significantly down-regulated in *L. marina* on the CeMM diet (Fig 6C).

The reduction of protein synthesis prolongs the lifespan of *C. elegans* (Syntichaki et al., 2007; Mittal et al., 2017; Zhou et al., 2021). The general decreases in protein synthesis can reduces the cellular load of erroneously synthesized polypeptides, cellular energy savings, coupled with reduced production of toxic metabolic derivatives in *C. elegans* (Hipkiss, 2007; Syntichaki et al., 2007). We found that RNA transcription and protein generation related genes, including ribosome biogenesis, ribosome, RNA transcription, splicing, protein processing in transport associated genes, were significantly down-regulated when *C. elegans* feeds on CeMM (Figure 5), which indicated that the CeMM diet might extend longevity of *C. elegan* via decreased protein synthesis related processes. Notably, the above transcription and protein synthesis related pathway genes were not enriched in *L. marina* on the CeMM diet.

A previous report showed that the increased levels of bile acids in the long-lived mice upregulate xenobiotic detoxification genes through the farnesoid X receptor (Amador-Noguez et al., 2007). Consistent with this, we found that three drug and cytochrome P450 xenobiotic detoxification pathway genes were significantly up-regulated in *C. elegan* on CeMM (Figure 6A). Our data suggested that the CeMM diet might extend the longevity of *C. elegans* on CeMM via up-regulation of the xenobiotic detoxification pathway. Notably, three drug and cytochrome P450 xenobiotic detoxification pathway related genes were significantly down-regulated in *L. marina* on the CeMM diet (Figure 5A). Although one P450 pathway genes, such as *gst-7 gst-8*, were significantly up-regulated in *L. marina* on the CeMM diet (Supplementary Figure 2A), we supposed that these genes might not involve longevity regulation. Given xenobiotic detoxification pathway genes were significantly contracted in the *L. marina* genome compared to *C. elegans* (Xie et al. 2020), it might indicate that a large number of xenobiotic detoxification related genes need to work together to promote animal lifespan.

### CeMM Attenuates the Development of Both *C. elegans* and *L. marina*

Given CeMM diet attenuate the development of both *C. elegans* and *L. marina*, the shared pathway in *L. marina* and *C. elegans* may play essential roles in attenuating their development in CeMM environments (Figure 6, Supplementary Figure 2). Further studies are required to delineate which gene play a major role.

We found that a series of RNA and protein processing related genes were significantly down-regulated in *C. elegans* under the CeMM diet (Figure 4). Breakdown of RNA assemble and transport can have catastrophic consequences like impaired RNA localization and distal protein synthesis that correlate with significant functional impairments such as neuronal growth defect (Fallini et al., 2016; Fernandopulle et al., 2021). RNAi of *eif-6*, which encodes the ortholog of vertebrate anti-association factor eIF6, slow down the growth of *C. elegans* (Fraser et al., 2000). RNAi of some structural constituents of ribosomes genes like *rps-12* and *rps-26* result in larval developmental delay or arrest (Melo and Ruvkun, 2012). Similarly, RNAi of *rpc-1*, which contribute to RNA polymerase III activity, *hel-1*, which predicted to enable RNA binding activity and RNA helicase activity, lead to slow growth (Kamath et al., 2003; Eki et al., 2007). The initiation factors are significant for protein transcription and regulation, of which downregulation could decrease the translation for most mRNAs under starvation or stress conditions (Sonenberg and Hinnebusch, 2009). Methionyl-tRNA synthetase (MRS) down-regulation result in DNA damage and further lead to functional defects (Kim et al., 2019). RNAi of protein processing in endoplasmic related genes *hsp-1* could result in larval developmental delay or arrest phenotypes (Melo and Ruvkun, 2012). RNAi knocks down *oxa-1*, which is involved in mitochondrial cytochrome C oxidase assembly and protein insertion into the mitochondrial inner membrane from the matrix, exhibits increased longevity, a reduced developmental rate, and increased resistance to oxidative stress (Maxwell et al., 2013). Given development requires building blocks like proteins to support cell growth and differentiation, our data suggest that the CeMM diet might attenuate the development of *C. elegans* on CeMM via down-regulation of protein synthesis related genes.

Unlike *C. elegans*, RNA and protein synthesis processing related genes were not enriched in *L. marina* on the CeMM diet. Of note, we found that the synthesis of unsaturated fatty acids pathway genes was significantly decreased in *L. marina* on the CeMM diet (Figure 6C). Unsaturated fatty acids have been reported to participate in numerous biological processes, like inflammation and immune response (Glass and Olefsky, 2012), neuro-reception (Battista et al., 2012), and reproduction (Hoang et al., 2013). *fat-2*, *fat-6*, and *fat-7* mutants, which have defects to synthesize 18- and 20-carbon unsaturation fatty acid, grow slowly compared to the wild-type (Watts, 2002; Brock et al., 2007; Vrablik and Watts, 2013). The depletion of *elo-1* and *elo-2* results in decreased PUFA, slow growth, and disrupted reproduction (Kniazeva et al., 2003). RNAi treatment of *elo-3* in both the *ire-1* and *atf-6* mutants background exhibited synthetic growth defects in *C. elegans* (Sakaki et al., 2012). In *L. marina*, we found that the expression of *elo-3* was significantly repressed under the CeMM diet. The data indicate that the CeMM diet might attenuate the development of *L. marina* on CeMM via down-regulation of unsaturated fatty acid synthesis genes.

## CONCLUSION

In this report, we found that the CeMM diet attenuated development of *L. marina* but did not promote its longevity, suggesting that the same chemically defined diet exhibit the same function on development attenuation but different role in longevity regulation in the two nematodes, since CeMM diet slow down the growth and extend the lifespan of *C. elegans* (Zhang et al. 2015). The present study will provide an essential foundation and basis for identifying the key genes and genetic pathways required for longevity and development, and further studies could manipulate the chemical composition of CeMM to identify how dietary components and their amount regulate animal development and longevity. Further genetic analysis of crucial gene(s) of longevity and growth in *C. elegans* and *L. marina* will provide imperative insights into the physiological changes of marine and terrestrial organisms in response to changing nutritional environments in the context of global climate change.

## DATA AVAILABILITY STATEMENT

The datasets generated for this study can be found in NCBI, and the BioProject ID: PRJNA801608

## AUTHOR CONTRIBUTIONS

PS and LZ conceived and designed the experiments. PS carried out most of the experiments, analyzed the data, and wrote the manuscript. XC was involved in sampling and data analysis.

LZ edited the manuscript and supervised the project. All authors read and approved the final manuscript.

## FUNDING

This work was funded by the National Key R and D Program of China [No. 2018YFD0901301]; “Talents from overseas Program, IOCAS” of the Chinese Academy of Sciences; “Qingdao Innovation Leadership Program” [Grant 16-8-3-19-zhc]; and Key deployment project of Centre for Ocean Mega-Research of Science, Chinese Academy of Sciences.

## ACKNOWLEDGMENTS

We are grateful to all members of the LZ laboratory for their helpful discussions.

## Notes

### Competing Interest Statement

The authors have declared no competing interest.

